# Repeated within-session rule switching in the marmoset: a paradigm for tracking the dynamics of cognitive flexibility

**DOI:** 10.64898/2026.07.31.740852

**Authors:** Marium H. Alvi, Ryley P. Nathaniel, Karmen Rai, Liya Ma

## Abstract

Adapting behaviour when reward contingencies change is a core function of cognitive control, but the underlying trial-by-trial computations are hard to observe when tasks cue each rule or allow only one switch per session. We developed the Feature-Rule Switching Task (FRST), in which common marmosets (*Callithrix jacchus*) repeatedly switch, without cues and under a fixed stimulus set, among the visual features that earn reward, inferring each switch from feedback alone. All four animals acquired the task within two days and sustained several switches per session across months of testing. A reinforcement-learning model with a learned weighting of stimulus dimensions best explained choices in every animal, outperforming complexity-matched perseveration controls. Learning-rate estimates fell within the human range, and in almost every session the animals weighted a stimulus dimension rather than individual features alone, with a dimensional commitment comparable in strength to that of humans. FRST thus provides a primate paradigm, amenable to laminar recording, for tracking the dynamics of cognitive flexibility.

## INTRODUCTION

Adapting one’s actions to changing contexts and contingencies is a fundamental component of cognitive control. For example, while driving, our attention shifts among priorities including monitoring the preceding cars, scanning the mirrors, reading road signs, and watching for pedestrians, based on the context of the road. Going beyond daily tasks, this process supports successful transition to adulthood (Moffitt et al., 2011), resilience against cognitive decline (Stern, 2012; Liu et al., 2024), and is disrupted across a range of neuropsychiatric conditions (Uddin, 2021; Grant and Chamberlain, 2023). Within the Research Domain Criteria (RDoC) framework, this ability is captured by goal updating and maintenance (Insel et al., 2010).

Often studied using rule-switching paradigms, goal updating involves multiple processes, including detecting changes in contingency, exploring alternative actions, and stabilizing newly effective strategies. Disentangling these processes requires experimental designs that allow updating to be observed as it unfolds over time. In many nonhuman primate studies, task-relevant mappings are explicitly instructed on each trial (Buschman and Miller, 2007; Kamigaki et al., 2009, 2012; Siegel et al., 2015). While these designs provide well-defined epochs for rule representation and action selection, they reduce the need for inference and exploration and primarily capture the implementation of known mappings. Conversely, feedback-based paradigms such as the intra- and extra-dimensional set-shifting task (IEDS) allow subjects to learn relevant stimulus features through trial and error (Sahakian et al., 1988; Owen et al., 1991; Cools et al., 2001). A hallmark of IEDS performance is the formation of an attentional set—a bias toward the previously trained and relevant stimulus dimension—which facilitates further shifts within that dimension (intra-dimensional) but impairs shifts to a new and never reinforced dimension (extra-dimensional). However, in animal models, the continency switches in IEDS are commonly performed across sessions, given the task difficulty due to the introduction of new stimuli. Thus, it does not permit the isolation of neuronal and ensemble dynamics of trial-by-trial goal updating from confounds related to time-on-task and motivation (Cowley et al., 2020; Leathers and Olson, 2012, Hyman et al., 2012).

A complementary line of work is the conceptual set-shifting and rule-switching tasks developed for macaques and humans (Moore et al., 2005; Ebitz et al., 2020; Goudar et al., 2024), which requires choosing and changing the target feature repeatedly within a session, based on feedback only. Because no new exemplars are introduced at a switch, these tasks do not require the formation of an attentional set. Instead, they expose the trial-by-trial dynamics of detecting a change, exploring alternatives, and re-stabilizing a rule, and they have proven well suited to latent-state modelling that separates rule-based from exploratory choices (Ebitz et al., 2020) and to quantitative comparison of rule-learning strategies across species (Goudar et al., 2024).

Cognitive control in the common marmoset (*Callithrix* jacchus) has so far been characterized almost entirely with reversal learning and IEDS (Roberts et al., 1988; Dias et al., 1996; Clarke et al., 2005; LaClair et al., 2019)—the same tests used to assay human executive function. This shared tradition makes human set-shifting data a natural reference point for marmoset cognition. Meanwhile, the within-session, recording-ready rule-switching paradigm now common in macaque and human work has not yet been established in the marmoset. For the neurophysiological investigation of frontoparietal circuits underlying goal updating, the common marmoset is a particularly well-suited. Unlike macaques, marmosets have a lissencephalic cortex, enabling simultaneous laminar recordings across lateral prefrontal and posterior parietal cortex (Johnston et al., 2019) — two regions central to cognitive control (Seeley et al., 2007). To leverage this advantage, however, a task is needed that produces repeated, within-session goal-updating behaviour under stable stimulus conditions.

To address these limitations, we developed a feedback-driven Feature-Rule Switching Task (FRST) for marmosets. In this task, animals learn to select the compound stimulus containing the target feature—a colour or a shape—based on trial-by-trial feedback. The same set of features is used throughout the session, and the rewarded feature changes repeatedly once performance criteria are reached. This design enables multiple within-session switches under stable stimulus conditions, allowing dissociation of error detection, exploration, and stabilization phases. In structure, FRST adapts the conceptual set-shifting and rule-switching tasks developed for macaques and humans (Moore et al., 2005; Ebitz et al., 2020; Goudar et al., 2024) to the marmoset; to our knowledge it is the first within-session, uncued rule-switching task established in this species, and it is built to yield the repeated switch events and stable stimulus conditions required by single unit recordings.

Additionally, goal updating and maintenance on feedback-based rule-switching tasks can be quantified as the learning rate and choice determinism in a feature reinforcement learning (fRL) model (Yearsley et al., 2021; Talwar et al., 2024). These parameters will guide the search for neuronal, ensemble and network-level mechanisms that support goal updating and maintenance. Additionally, similarity in these parameters to human performance can be used to gauge the translational potential of an animal model (Redish et al., 2021; Yamamori et al., 2023; Neville et al., 2024).

In the present study, we show that marmosets can perform rapid, repeated feature-rule switches within a single session and that their behavior is well described by an attention-augmented RL model whose learning rate and dimensional commitment fell within the human range for the IEDS task that leads the marmoset cognitive flexibility literature (Talwar et al., 2024). Across nearly every session the animals organised their choices around a stimulus dimension, even though FRST can be solved by tracking individual features. These findings establish FRST as a translational paradigm for quantifying goal updating and lay the groundwork for future investigation of the neural mechanisms supporting flexible behaviour.

## RESULTS

We trained marmosets to perform the Feature-Rule Switching Task (FRST) using a home cage-attached touchscreen training box. We started by pretraining them on Simple Discrimination on shape and colour stimuli, with every feature serving in turn as the rewarded target and as the foil (**Figure 1A**). Then in the full version of FRST we presented them with compound stimuli (i.e. coloured shapes) for the first time (**Figure 1B**). Marmosets must touch the stimulus containing the target feature (e.g. heart) to obtain a reward, ignoring the other features. Once they reached 80% accuracy, the target feature switched without any cue—alternatingly to a feature within the same dimension (e.g. heart to star) or cross dimensions (e.g. heart to yellow)—so that flexibility could be tracked continuously within a session (**Figure 1C**). After sufficient data had been collected, the task was repeated with an entirely new set of shapes and colours (**Figure 1D**), without pretraining, in order to examine whether they could transfer their experience to unfamiliar features.

**Figure 1.**
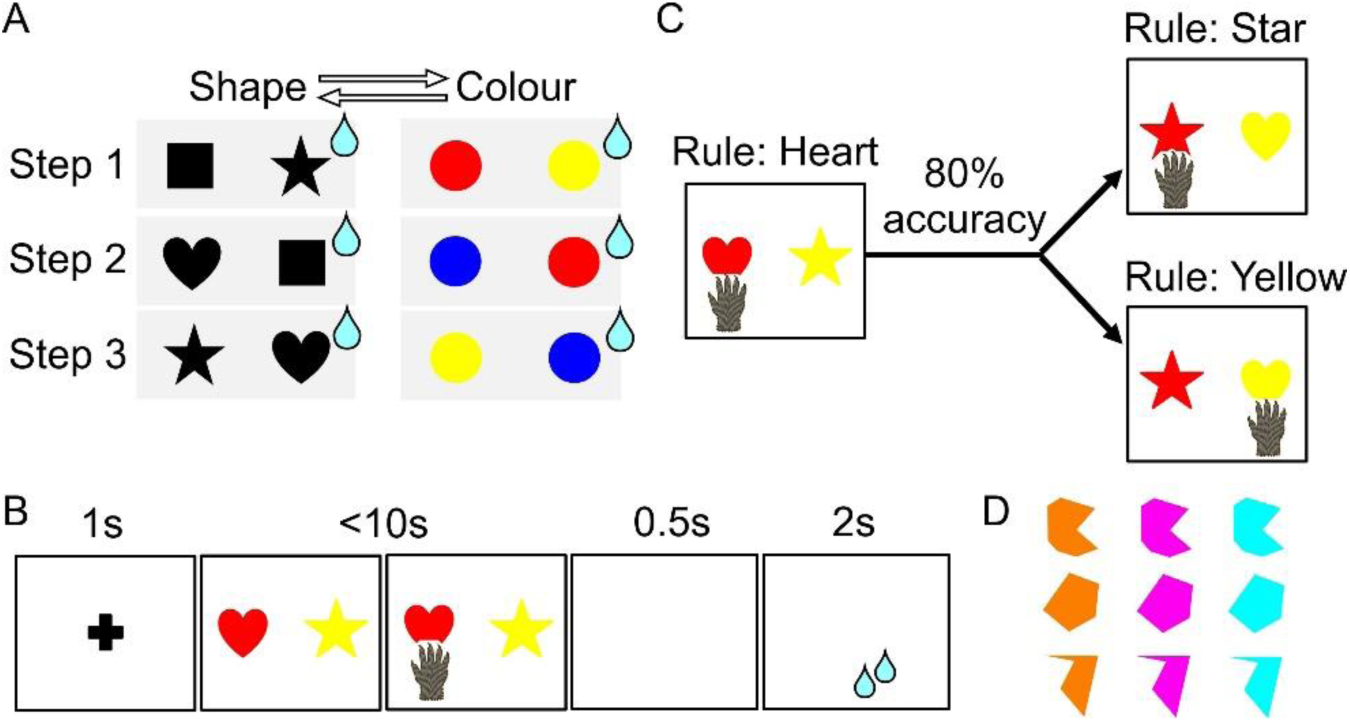
Schematics for training steps and the FRST. **A)** Initial shape and colour training for the original stimulus set. Each shape and colour acted as a target and a foil. **B)** Within-dimensional switches are when the target feature changes within a dimension, e.g. heart to star, upon meeting the 80% correct response criterion. An cross-dimensional switch is when the target feature changes across dimensions (e.g. heart to yellow). **C)** Rule switching touchscreen task. **D)** New Stimuli set shapes and colours.

### Marmosets learned to consistently shift target feature multiple times within session

By Day 2 of FRST training, all 4 marmosets achieved 2 or more rule switches within session. These included switches between features both within and cross dimensions. In an example session (**Figure 2**), in 50 trials, Marmoset M reached criterion by choosing the heart-shaped stimuli while ignoring the colour or location (**Figure 2**, blue shade; red horizontal bar highlights at-criterion performance). At this point, the rule switched to ‘choosing red-coloured stimulus’ (**Figure 2**, red shade). The animal again reached criterion, in approximately 20 trials. The third and fourth target features were ‘yellow’ and ‘star’ (**Figure 2**, yellow and purple shades), respectively, before it switched back to ‘heart’ as the fifth target. In this session, Marmoset M completed 6 switches in a total of 220 trials.

**Figure 2.**
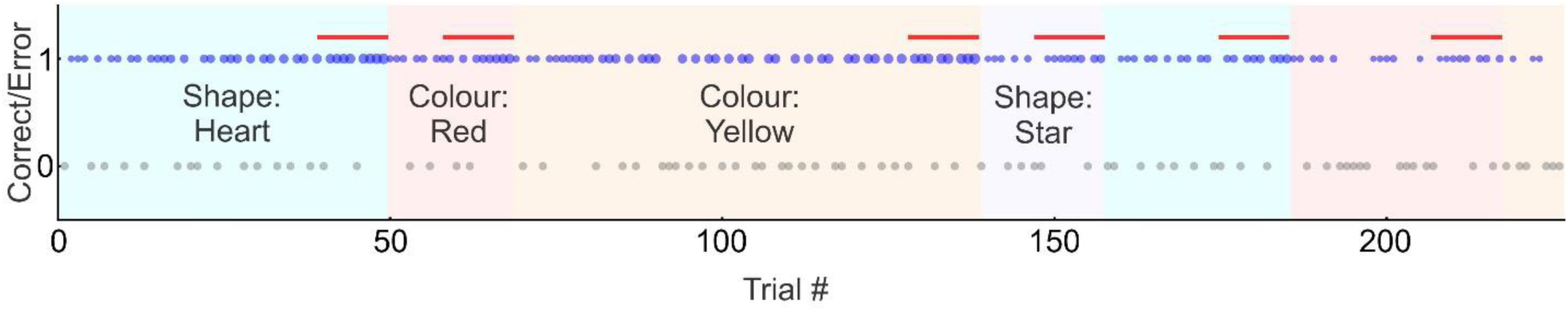
Intra- and extra-dimensional shifts within an example session. Shaded background regions denote the correct feature rule. Blue dots denote correct choices, with sizes increasing with the length of the successful streak. Gray dots represent errors. Red horizontal bars above the trials indicate the trial block that reached the criterion.

Overall, using the first feature set, the 4 marmosets completed 73 sessions in total with an average of 3.60 switches per session (median = 4, range = 1-9) (**Figure 3A**). Given this experience, using the new set of features, marmosets entered the full version of the task without additional Simple Discrimination training and started switching targets multiple times on Day 1 (**Figure 3B**). In a total of 90 sessions, they completed an average of 3.61 switches per session (median = 4, range = 2-9) (**Figure 3B**).

**Figure 3.**
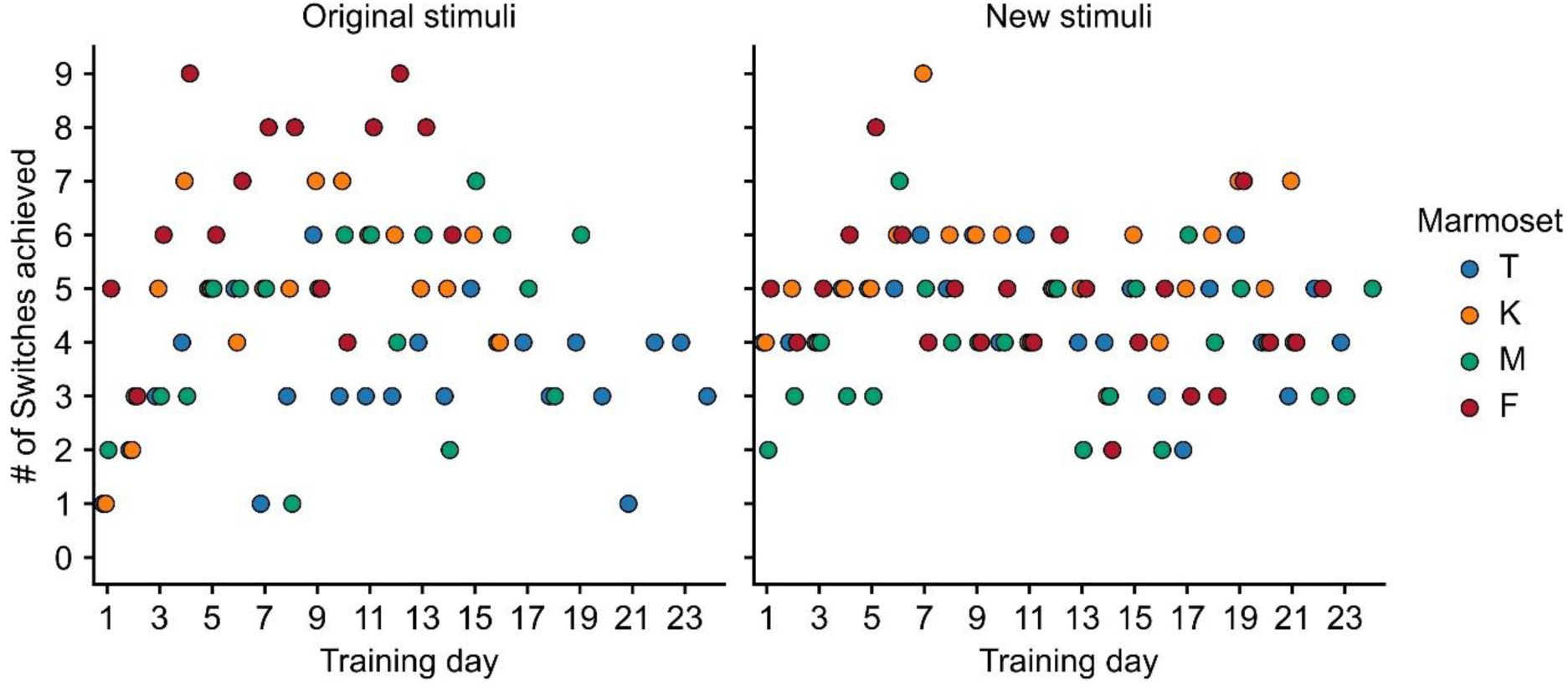
Number of rule switches achieved across training days. Colours denote individual animals. Left panel shows performance with the first set of features and the right panel for the second feature set.

We then examined their performance before and after switches between features within vs across dimensions. We defined a ‘switch block’ as a block of 14 trials including 7 trials before and after the switching point, respectively. As expected, switches were marked by high accuracy at 80% just before and a significant drop in performance in all animals (range: 47.4% to 55.6%) for the original (**Figure 4**) and the new features (**Figure 5**). A mixed-model ANOVA revealed a significant main effect of pre-versus post-switch performance for both the original feature set (*F*_1,3_ = 997.65, *p* = 7.0 × 10⁻⁵) and the new feature set (*F*_1,3_ = 1078.23, *p* = 6.2 × 10⁻⁵), confirming a robust reduction in accuracy immediately after feature-rule changes. We also found a main effect of switch type, *F*_1,3_ = 11.77, *p* = .042, indicating that post-switch accuracy differed between within- and cross-dimension switches, with slightly higher accuracy observed following the latter type of switches. This is expected, because when the target feature switched within dimensions (e.g. red to yellow), continued choice of the previous feature was now wrong every time (0%); but when the switch crossed dimensions (e.g. red to heart), choosing the previous feature was now still correct when the pre- and post-switch target features co-localize (50%, e.g. compound stimulus is red heart). There was no main effect of previous rule (*F*_1,3_ = 2.02, *p* = .25) or feature set (*F*_1,3_ = 0.06, *p* = .83).

**Figure 4.**
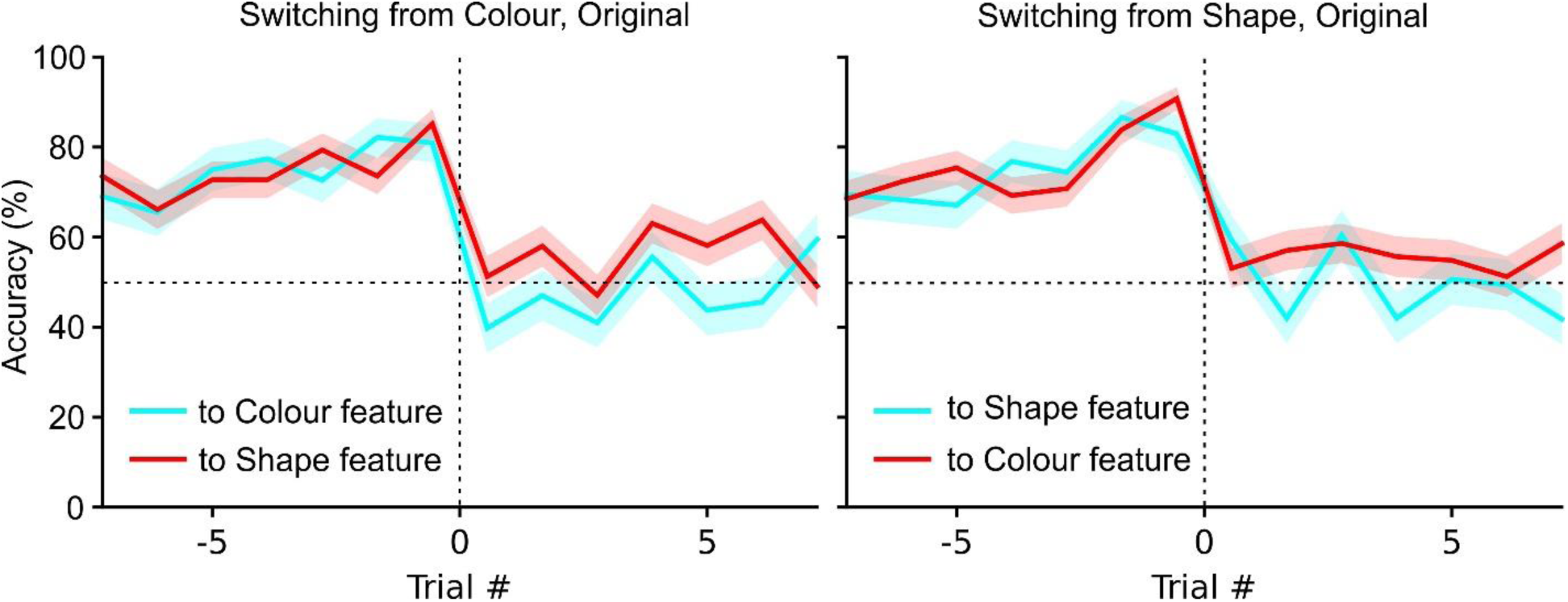
Performance around target switching with the first set of features, from: **A)** a colour feature, or **B)** a shape feature. Cyan lines indicate switching within a dimension, red indicate switching across dimensions.

**Figure 5.**
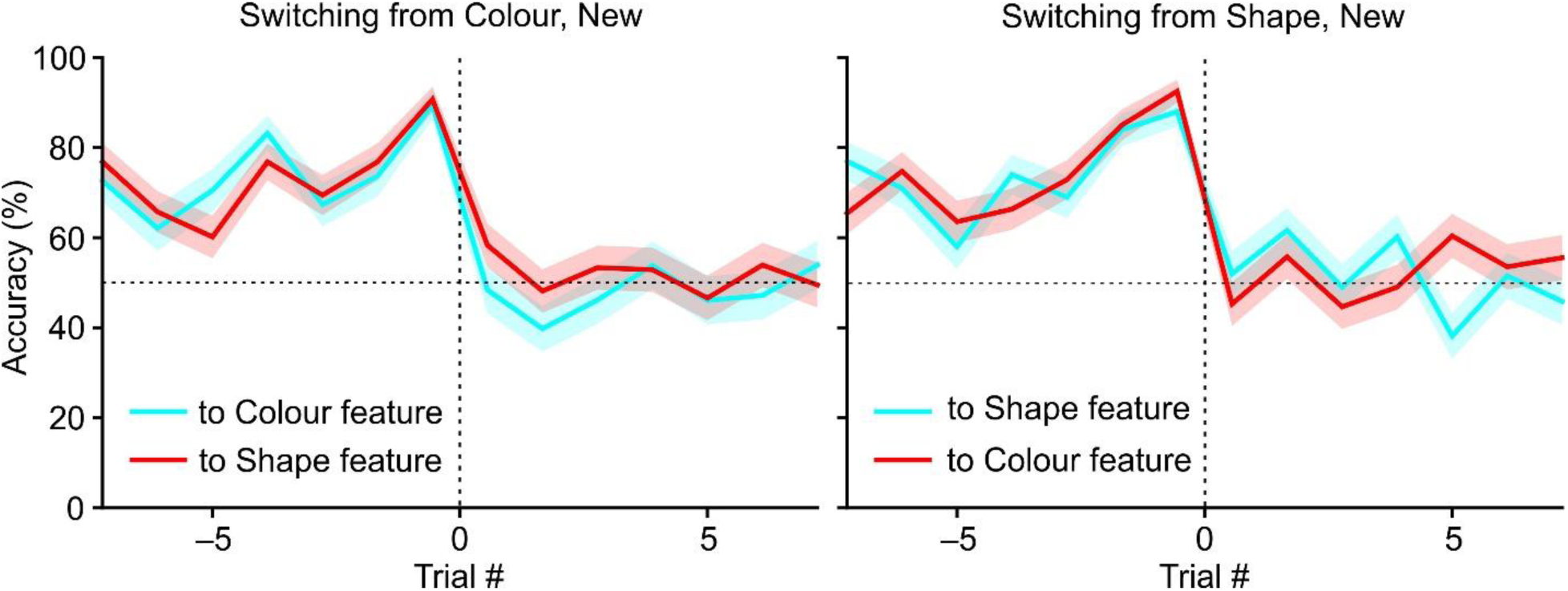
Performance around target switching with the second set of features, from: **A)** a colour feature, or **B)** a shape feature. Cyan lines indicate switching within a dimension, red indicate switching across dimensions.

### Perseverative errors had longer reaction times

Having found reliable performance just before each switch, we next asked whether their errors immediately after a switch reflected choices guided by the previous, now incorrect, rule. When marmosets made a wrong choice, the same stimuli were shown again on the following trial as a chance to ‘try again’. Animals chose correctly on this second chance on average 81.8% of the time, within the first 10 post-switch trials (**Figure 6**). This left 18.2% of the time when marmosets chose the same, incorrect stimulus twice in a row. These perseverative errors—errors that occurred after an error and involved the same wrong choice, had significantly *longer* reaction times (RTs) compared to correct choices made on such second-chance, for 3 of the 4 marmosets (**Figure 6**) with both the Original feature set (2-way ANOVA, overall *F*_1,3_ = 10.95, p = .045; post hoc Tukey’s test: p <= .001 for Marmosets F and M, p = .042 for K; but p = .861 for T), and the New feature set (p < = .001 for F, K, and M; but p = .623 for T). There was no significant main effect of feature set (*F*_1,3_ = 2.04, p = .248), and no significant interaction between correct vs error trial type and feature set (*F*_1,3_ = 0.53, p = .520). By contrast, following correct trials, RTs did not differ between correct and error responses for any animal in either feature set (all p >= .591). The extra-long RTs suggest that perseverative errors were not simply rapid or impulsive repeats. While we defined these errors as perseveration, they also reflected a true adherence to the previous rule, as the animals knowingly rather than impulsively choose the same unrewarding choice again, as if seeking confirmation that the rule had indeed switched.

**Figure 6.**
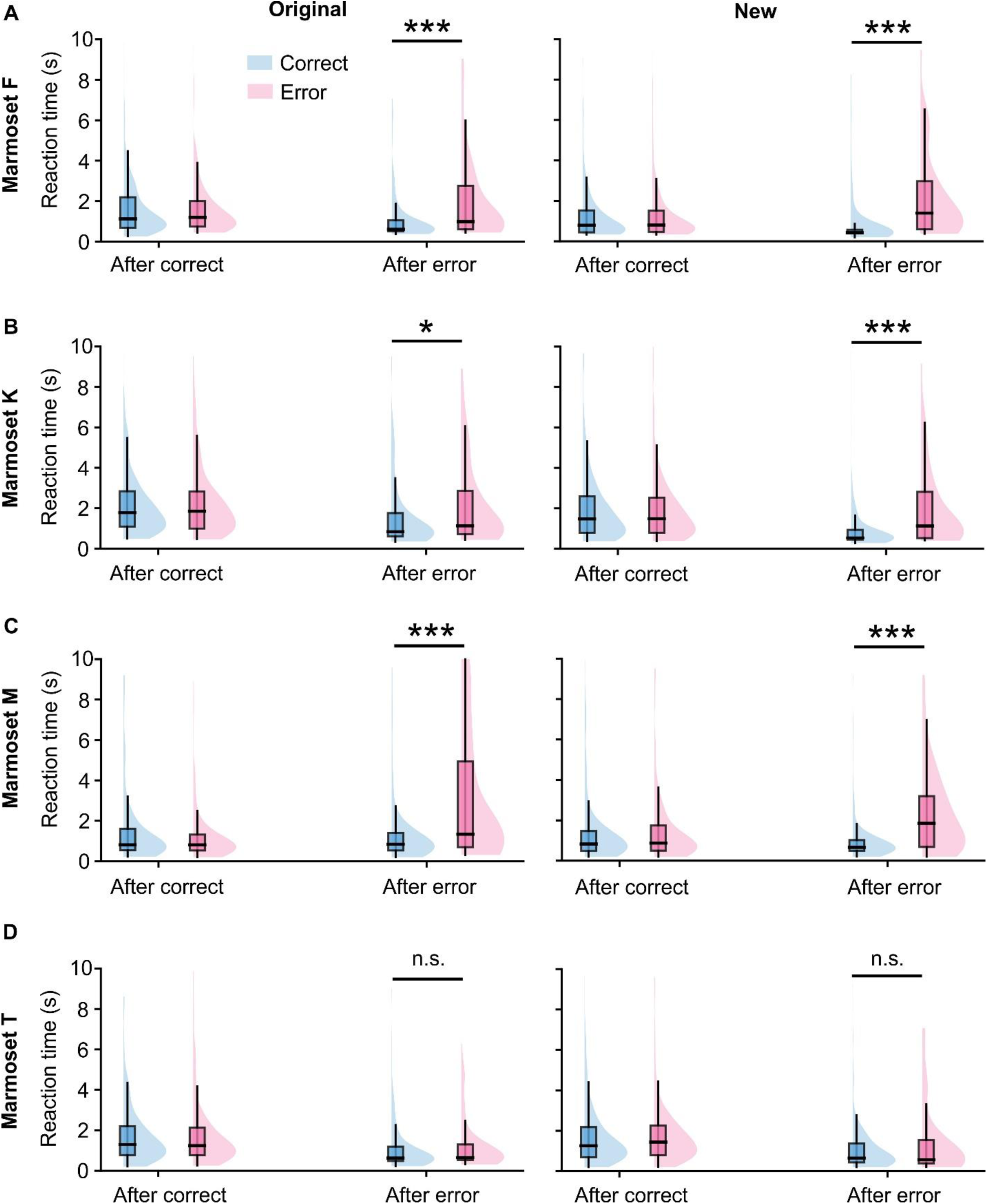
Reaction times (RTs) of correct choices and errors in the first 10 trials following rule switches, displayed separately for the original stimulus set (left panels) and the new stimulus set (right panels). Within each panel, RT distributions are separated by the outcome of the preceding trial (After Correct vs After Error) and the current choice (Correct: blue; Error: pink). *–p < 0.05; ***–p <0.001.

### An attention-augmented reinforcement-learning model best accounts for trial-by-trial errors

To reveal the learning and attentional processes that generate the marmosets’ task performance, and to compare them to similar processes underlying human performance on the well-established IEDS task, we fit three reinforcement learning models to the animals’ choices trial by trial. These include Feature Reinforcement Learning (fRL), Combined Attention-Modulated Feature Reinforcement Learning (Ca-fRL), and Separate Attention-Modulated Feature Reinforcement Learning (Sa-fRL) (Talwar et al., 2024). All three model computes stimulus values based on their features, then uses a softmax function to produce the choice and updates feature and/or attention weights via prediction errors. The fRL model updates only feature weights with no consideration for attention to the Colour vs Shape dimensions. Ca-fRL model extends the fRL by introducing a weight for attention to the previously relevant dimension; and the Sa-fRL further extends the Ca-fRL by introducing a second learning rate, that governs the updates to the dimension attentional weights (Talwar et al., 2024).

To compare the models while penalising for the number of free parameters, we computed the Bayesian Information Criterion (BIC) for each model, animal, and stimulus set (**Table 5**). Ca-fRL consistently yielded the lowest BIC in every animal and for both stimulus sets (8 of 8 cases), while the feature-only FRL model always yielded the highest BIC. This uniform pattern indicates that adding a dimensional-attention layer improves the penalised fit over feature-only reinforcement learning, and that the second, separate dimension learning rate introduced in SafRL was not justified. Because all three models share the same feature-learning core, this comparison suggests that dimensional attention played a role in addition to feature-based learning.

**Table 5.** Bayesian Information Criterion (BIC) values by model, stimulus set, and animal (O: Original and N: New feature sets). Lower values indicate a better balance of fit and parsimony. Ca-fRL (bold) yielded the lowest BIC in all four animals and both stimulus sets.

| ID | Original features |  |  |  |  | New features |  |  |  |  |
| --- | --- | --- | --- | --- | --- | --- | --- | --- | --- | --- |
|  | FRL | Ca-FRL | Sa-FRL | PFRL3 | PFRL4 | FRL | Ca-FRL | Sa-FRL | PFRL3 | PFRL4 |
| M | 492.91 | **466.73** | 475.32 | 480.69 | 502.03 | 308.42 | **279.07** | 289.07 | 303.61 | 324.79 |
| T | 382.53 | **353.80** | 363.98 | 380.16 | 392.15 | 396.78 | **366.92** | 367.16 | 382.48 | 400.99 |
| K | 481.94 | **450.47** | 457.24 | 479.99 | 509.46 | 455.37 | **422.33** | 428.95 | 444.91 | 472.99 |
| F | 640.82 | **583.10** | 599.52 | 631.50 | 660.32 | 599.70 | **552.51** | 560.72 | 574.74 | 583.02 |

We then asked whether the better fitting of the 3-parameter Ca-fRL compared to the 2-parameter fRL could be explained by the addition of a task-related, non-dimensional mechanism. Choice perseveration is a suitable choice, as it allows the test of whether the ‘attention to dimension’ could explained by a tendency to repeat recently chosen features. We therefore added two feature-RL models expanded with a feature-level choice-perseveration term (**Table 5**): a last-trial form with three free parameters, matched in number to Ca-fRL, and a decaying choice-kernel form with four free parameters, matched to Sa-fRL. Both perseveration models yielded higher BIC than Ca-fRL and Sa-fRL. Given the matched number of parameters, these differences cannot be attributed to the complexity penalty. That is, at matched complexity, dimensional attention provided a better account of marmoset choices than choice perseveration, and even the more flexible four-parameter perseveration model did not surpass the three-parameter Ca-FRL. Choice perseveration therefore does not account for the fit advantage of the attention-augmented models.

### The results of feature-RL model fitting with marmoset FRST performance closely resembled those from human IEDS data

Identifying the model that best captures marmoset behaviour lets us ask whether the computations underlying marmoset flexibility correspond to those of humans performing an analogous set-shifting task. We first checked that the models reproduced marmoset behaviour as well as they reproduce human behaviour, then compared the fitted parameters against the human estimates directly.

As a qualitative posterior-predictive check that complements the BIC comparison, we show the correlation between observed and model-predicted error rates for each marmoset and model (**Figure 7**, blue: original feature set, orange: new feature set). Remarkably, these plots are visually indistinguishable from those created from human IEDS performance in Figure 2A of Talwar *et al* (2024). Pearson correlation coefficients and significance levels are shown in each panel. The fRL model failed to capture behavioural error patterns for three of the four animals (Marmosets M, F, and K, all p ≥ .09). For Marmoset T, fRL correlated with performance for the new feature set (r = .53, p = .02) but not for the original set (r = .30, p= .16), indicating that a single learning rate reinforcement learning model does not reliably account for trial-level error dynamics. In contrast, Ca-fRL produced uniformly strong correlations across all animals and both feature sets (all r ≥ .84, p < .001) and Sa-fRL showed similarly high performance (all r ≥ .86, p< .001), with correlations ranging from .86 to .92 for the new stimuli and .88 to .98 for the original stimuli. These posterior-predictive checks mirror the BIC comparison: the feature-only model fails to track trial-level error dynamics in most animals, whereas both attention-augmented models reproduce them closely. As with BIC, the two attention-augmented models are difficult to separate on this measure (Ca-fRL and Sa-fRL differ negligibly), so for the parameter analyses below we adopt Ca-fRL, the more parsimonious of the two and consistently with the lower BIC.

**Figure 7.**
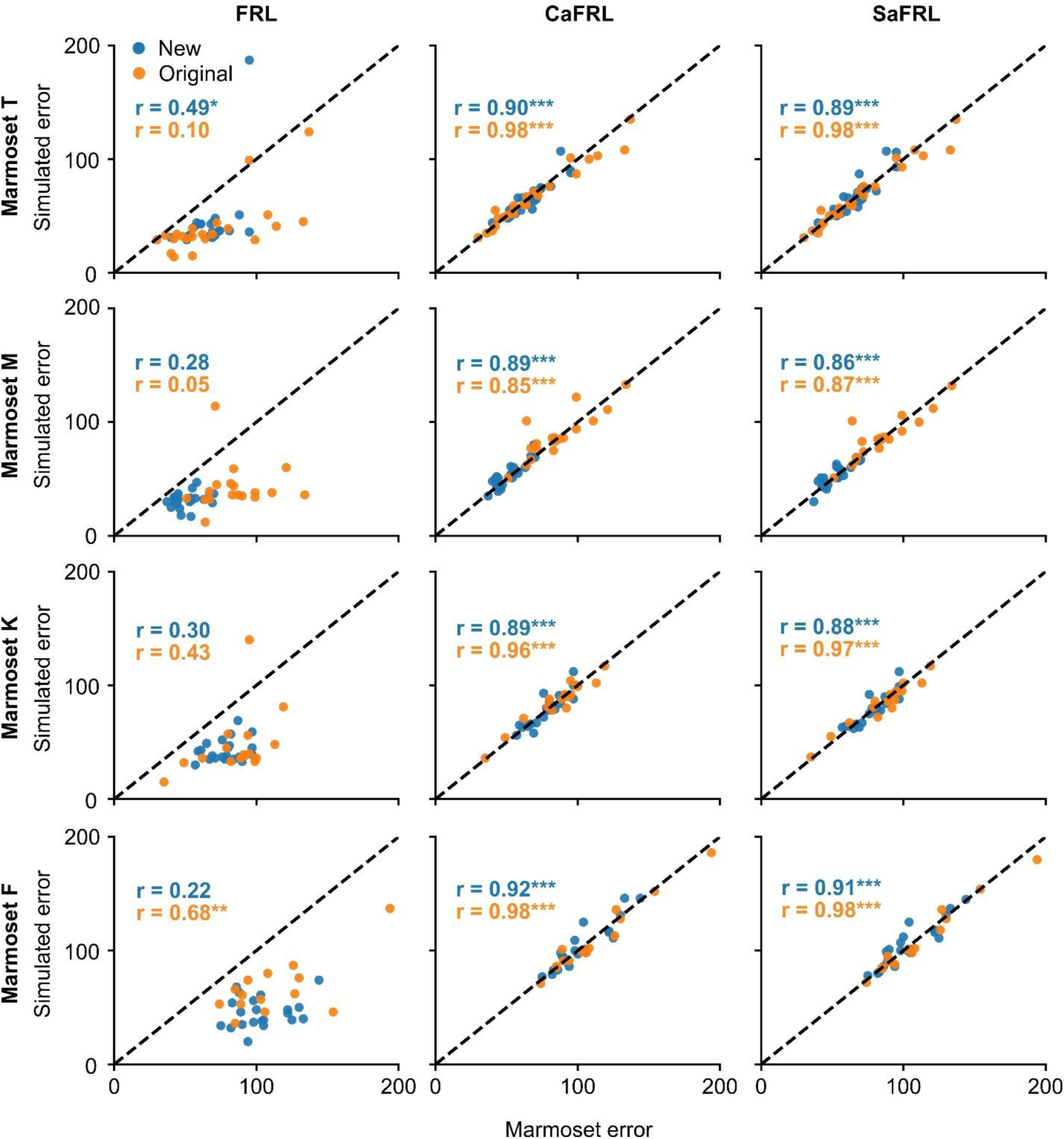
Trial by trial error correlations for FRL, CaFRL and SaFRL models across four marmosets. Each panel plots observed marmoset error counts against model predicted errors for the new stimuli set trials (blue circles) and the original stimuli set (orange circles). Rows correspond to individual animals (Marmoset T, M, K and F) and columns to the three models (FRL, CaFRL, SaFRL). Dashed diagonal lines indicate perfect prediction. Pearson’s r and p-values for each condition are displayed within the legends of each panel.

We next compared the fitted Ca-fRL parameters — learning rate (α), choice determinism (β), and dimensional weighting — against the corresponding human values reported by Talwar et al. (2024) for the intra-/extra-dimensional set-shifting (IEDS) task (**Figure 8**). Because our marmosets acquired and sustained FRST proficiently, we asked whether their per-session estimates fell within the range of human performance, taking the human mean ± one standard deviation as the reference range for each parameter.

**Figure 8.**
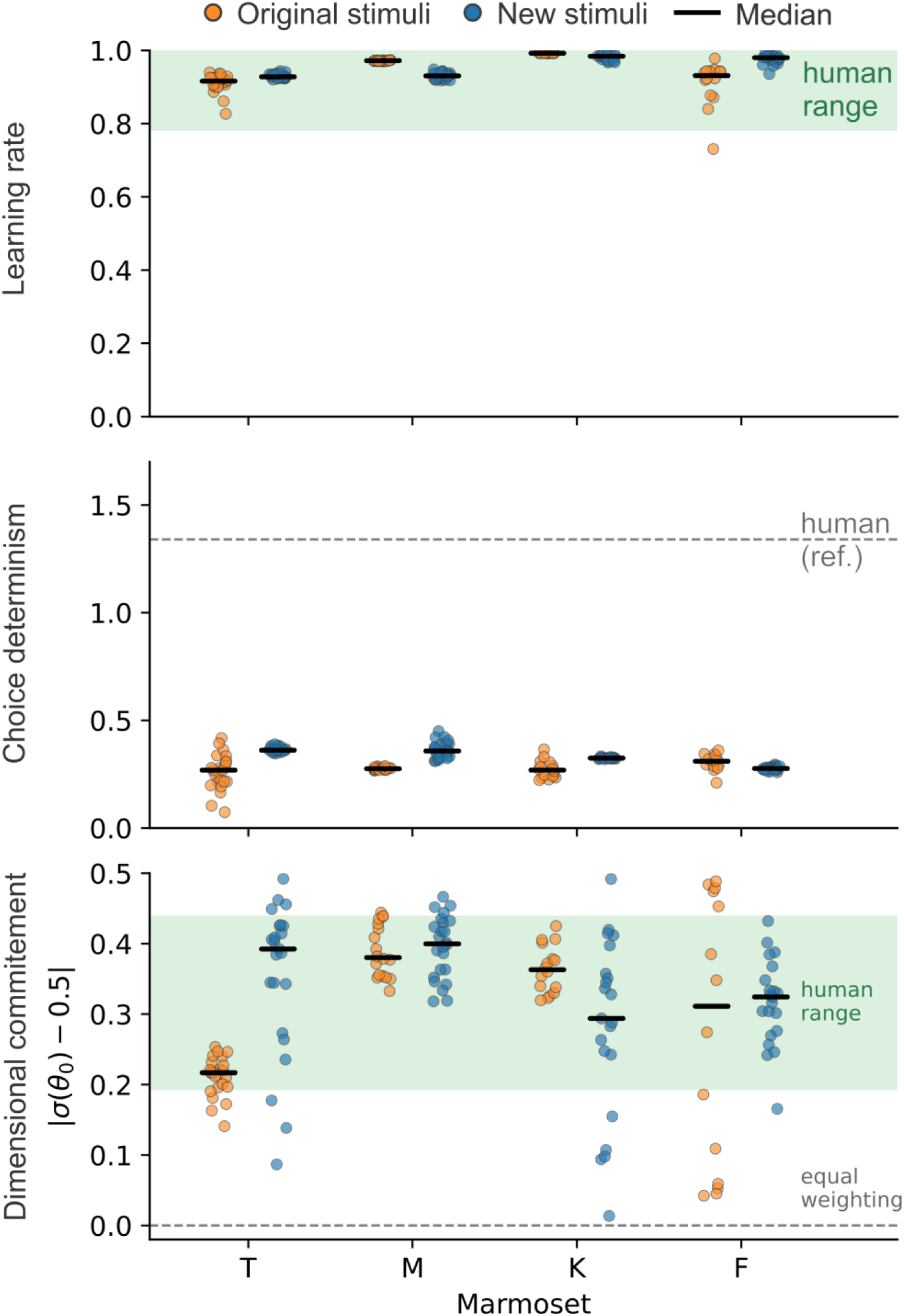
Ca-fRL parameters for each marmoset (T, M, K, F), with every session shown as a dot; orange = original feature set, blue = new feature set, and the black bar marks the per-cell median. Top: learning rate, with the human range (mean ± 1 SD; Talwar et al. 2024) shaded. Middle: choice determinism, with the human mean shown only as a reference (dashed line). Bottom: dimensional commitment, |σ(θ_0_) − 0.5|, where 0 is equal weighting of the two dimensions and 0.5 is all weight on one dimension; the shaded human range is obtained by passing the human dimension-primacy estimate through the same transform. Commitment is a magnitude and does not distinguish colour from shape, so the human-range overlap reflects equivalence of commitment strength, not of which dimension was attended. Per-session dots are shown rather than a mean because averaging signed weights cancels the two dimensions.

To test whether each animal’s estimates fell within the human range, we used two one-sided tests of equivalence (TOST) against the human values of Talwar et al (2024), with equivalence bounds set a priori at ±1 human SD and each animal’s sessions as its repeated measurements (Materials and Methods). Every animal’s mean learning rate (α) fell inside the human band [0.78–1.06] for both feature sets (original/new — T: 0.85/0.86; M: 0.95/0.88; K: 0.83/0.86; F: 0.79/0.90), and equivalence was confirmed in six of the eight animal × feature-set cases (all p < .05). The two cases that did not reach equivalence (K and F, original set) had means well inside the band but greater session-to-session variability across fewer sessions, so the test was inconclusive rather than indicating a value outside the human range; both reached equivalence on the new set. Marmoset learning rates therefore lie within the range reported for humans on IEDS.

Dimensional weighting is best summarised as commitment — how far the fitted weighting departs from equal weighting of the two dimensions, |σ(θ_0_) − 0.5|, where θ_0_ is the dimensional primacy parameter obtained from Ca-fRL, and a commitment value of 0 denotes equal weighting and 0.5 all weight on one dimension (**Figure 8**). Commitment was substantial across the dataset: 95% of sessions exceeded 0.1 and the median was 0.35, indicating that the animals organised their choices around a stimulus dimension even though FRST can be solved without attending to a dimension. Every animal’s commitment fell within the human range [0.19–0.44] obtained by passing the human dimension-primacy estimate through the same transform. Because commitment is a magnitude that does not distinguish colour from shape, it indexes how strongly, not which, dimension was favoured; the signed per-session primacy estimates (θ_0_) were weakly identified (large session-to-session variability), and we therefore neither interpret their direction nor compare them across feature sets.

Choice determinism (β) was consistently lower in marmosets (∼0.3) than in the human estimate (1.34) across all animals and both feature sets. Because inverse-temperature parameter scales inversely with the magnitude of the values it multiplies and because FRST and IEDS differ in reward coding and trial count, the absolute value of β is not directly comparable across the tasks. We therefore report choice determinism for completeness but did not test it for equivalence. It remains a possibility that a lower value of β indicates more exploratory choice in the marmosets compared to humans.

## DISCUSSION

We introduced the Feature-Rule Switching Task (FRST), a feedback-driven paradigm in which marmosets repeatedly update which stimulus feature they pursue, many times within a single session, without explicit cues and while the stimulus set remains fixed. All four marmosets learned to switch among familiar colour and shape features within two days of first encountering compound stimuli, and they continued to switch several times per session across months of testing. Each switch produced a sharp, transient drop in accuracy, after which performance recovered. When a wrong choice was repeated on the immediately following second-chance trial, these perseverative errors were slower than correct choices, suggesting continued adherence to the previous rule rather than impulsive or inattentive responding. Among the reinforcement-learning models we fit, an attention-augmented model (Ca-fRL) provided the best penalised fit (lowest BIC) in every animal and both stimulus sets, outperforming a feature-only model. Additionally, both its learning rate and dimensional commitment fell within the ranges reported for humans on a related set-shifting task. Together, these results establish FRST as a behavioural and computational paradigm for studying goal updating in a primate species well-suited to laminar recording in relevant cortical circuits.

FRST was designed to remove confounds that have limited the study of goal updating in nonhuman primates. Paradigms that cue the relevant rule on each trial isolate the implementation of an instructed mapping but bypass the inference and exploration that updating requires (Buschman and Miller, 2007; Kamigaki et al., 2009, 2012; Siegel et al., 2015). Feedback-based set-shifting tasks such as the IEDS restore that demand, but in animal models the contingency changes typically occur across sessions and are accompanied by new stimuli, so the dynamics of updating become intertwined with time-on-task, motivation, and stimulus novelty (Roberts et al., 1988; Dias et al., 1996; Clarke et al., 2005; Leathers and Olson, 2012; LaClair et al., 2019; Cowley et al., 2020). By holding the stimulus set fixed and driving many uncued switches within a session, FRST separates the moment the contingency changes from the exploration and re-stabilization that follow, and it yields the repeated switch events and trial counts required by intracortical recordings. Because the marmoset cortex is lissencephalic, lateral prefrontal and posterior parietal cortex are both accessible to simultaneous laminar recording (Johnston et al., 2019).

Cognitive control in the marmosets has been assessed almost exclusively with reversal-learning and intra-/extra-dimensional set-shifting tasks. The within-session, repeatedly-switching rule design has proven productive in macaques and humans (Moore et al., 2005; Ebitz et al., 2020; Goudar et al., 2024), in which the active rule changes several times per session and must be continually re-inferred. This paradigm has not, to our knowledge, been available in the marmoset. FRST imports that design, extending a paradigm already shown to support latent-state modelling of exploration (Ebitz et al., 2020) and cross-species comparison of rule-learning strategies (Goudar et al., 2024) to the marmosets.

The slowing of repeated errors also offers a behavioural window onto how the marmoset brain represented the task. Had these errors reflected lapses or impulsive responding, they should have been as fast as or faster than correct choices. Instead, they were slower, consistent with the animal deliberately selecting the previously rewarded feature while that feature was being extinguished. This pattern fits the broader view that updating proceeds through a detectable sequence of error detection, exploration, and re-stabilization (Ebitz et al., 2020; Goudar et al., 2024), and it provides a behavioural marker that can later be aligned with neural signatures of error monitoring (Mansouri et al., 2006). In macaques and humans, the same family of tasks resolves into latent rule-based and exploratory states (Ebitz et al., 2020), and slower, perseverative responding after a switch, together with reduced sensitivity to negative feedback and more variable exploration, distinguishes rule-learning in macaques from humans (Goudar et al., 2024). The perseverative slowing we observe provides a marmoset entry point into this framework and motivates a direct test of where marmosets fall along these axes relative to macaques and humans.

Furthermore, we asked how the Ca-fRL parameter estimates from FRST relate to those reported for humans performing the intra- and extra-dimensional set-shifting task (Talwar et al., 2024). Using two one-sided tests of equivalence, marmoset learning rates fell well within the human range. We use the human estimates as the reference frame through which marmoset cognitive control has most often been gauged, rather than as a matched control. Since marmosets have homologues to all major areas and networks of the human brain (Ghahremani et al., 2017; Majka et al., 2020), we interpret this overlap in learning rates as an indication of shared underlying computational processes (Redish et al., 2021; Yamamori et al., 2023; Neville et al., 2024).

Our BIC comparison shows that the dimensional-attention component earns its added parameters: Ca-fRL provided the best penalised fit in every animal and both stimulus sets, outperforming the feature-only model. However, to establish that dimensional structure is specifically responsible, requires ruling out equally flexible non-dimensional accounts. The most salient such alternative is choice perseveration: because repeated errors were slower rather than faster, a feature-RL model that simply favours recently chosen features—through choice perseveration or hysteresis—might in principle reproduce the CaFRL fit without dimensional modulation (Lau and Glimcher, 2005). We tested this possibility by adding to the basic fRL model a feature-level choice-perseveration term in two complexity-matched forms: a last-trial trace with the same number of free parameters as Ca-fRL, and a decaying choice kernel with the same number as SaFRL, and compared them under the same BIC criterion. Both perseveration models fit worse than Ca-fRL and Sa-fRL. At matched complexity, dimensional attention accounted for marmoset choices better than choice perseveration, and even the more flexible four-parameter perseveration model did not surpass the three-parameter Ca-fRL. Choice perseveration therefore does not offer a parsimonious alternative to dimensional modulation. This addresses the most likely non-dimensional confound, though not every conceivable one: other non-dimensional mechanisms, such as value forgetting or asymmetric learning from positive and negative feedback, remain to be examined, and full parameter- and model-recovery analyses would further secure the dimensional interpretation (Wilson & Collins, 2019).

The dimensional weighting itself is the clearest sign that the animals recruited stimulus dimensions. Expressed as commitment—the distance of the fitted weighting from equal weighting of the two dimensions—it was substantial in almost every session and comparable in strength to human values, even though FRST can be solved by tracking individual features without reference to any dimension. Notably, because commitment is a magnitude, it speaks to how strongly, not how appropriately, a dimension was weighted. That is, whether that weighting is held fixed within a session or is reallocated as the rewarded rule changes, and whether it follows the currently relevant dimension, are questions the current analyses cannot settle. Despite this limitation, as the best-supported model among all that we compared, the recruitment of dimensional structure under conditions that do not demand it is an unexpected finding.

Several limitations follow from the scope of this study. The sample comprised four male marmosets, which precludes analysis of individual variability and of sex as a biological variable (LaClair et al., 2019; Nephew et al., 2020), both deserve attention as the paradigm is extended. The present work is behavioural and computational, and the neural mechanisms we invoke remain to be tested directly. As noted, the human comparison is cross-task and qualitative. None of these undercuts the core behavioural result—that marmosets perform rapid, repeated, uncued feature-rule switches under stable stimuli; but each limit our capacity for drawing conclusions over the computational and cross-species comparisons.

FRST opens several lines of investigation. Its repeated within-session switches and fixed stimulus set suit it to simultaneous laminar recordings in lateral prefrontal and posterior parietal cortex (Johnston et al., 2019), to circuit perturbations that can target specific phases of updating (Jendritza et al., 2023; Shaw et al., 2026), and to linking the model-derived learning and attention parameters to neural activity. Because the same computational framework applies to human set-shifting (Yearsley et al., 2021; Talwar et al., 2024), FRST also offers a route for testing whether manipulations in the marmoset reproduce the parameter changes seen in human cognitive control and its disorders. By making goal updating observable as it unfolds, under conditions compatible with circuit-level measurement, FRST provides a foundation for a mechanistic and translational account of flexible behaviour.

## MATERIALS AND METHODS

### Subjects

Four wild-type, experimentally naïve male common marmosets (*Callithrix jacchus*) were used in this study. All were 2 to 3 years old, with weight ranging from 350–500g and remained healthy throughout the study. They were procured from SILABE (Strasbourg, France). Marmosets M and T were housed together in one cage, and Marmosets K and F were together in another. All procedures were carried out in accordance with the Canadian Council for Animal Care policies on care and use of laboratory animals, following Animal Use Protocol 2023-01 as approved by the Animal Care Committee of York University.

Marmosets were not restricted on food or liquid intake. The amount and variety of their last meal of the day were adjusted to match their needs, so no food would be left by next morning. Training was always conducted early in the morning before their first meal of the day.

### Apparatus

Two touchscreen training boxes (Neuronitek, London, Canada) were used in this study with in-house modifications to accommodate the following components. Each box was equipped with an Elo 10” touchscreen (Elo, Knoxville, TN, USA), a reward pump (NE510 Single Channel Programmable OEM, New Era Pump Systems Inc, Farmindale, NY, USA), and two Raspberry Pi HQ Cameras (Raspberry Pi Ltd., Cambridge, UK) at the top and left sides of the box. All components were controlled by a Raspberry Pi 4 Model B computer via custom-written software in Python. The reward was either dilute acacia gum or dissolved marshmallow fluff (Durkee Mower Inc., Lynn, MA, USA) chosen based on the animal’s preference.

### Pre-training

#### Touchscreen training

At this stage marmosets trained in 30-trial mini sessions. They progressed to the next step upon meeting criteria of 80% accuracy in 2 consecutive mini sessions. In each trial, a black filled circle extending the width of the screen was presented for 10s. If touched, the circle disappeared and a reward was delivered after a brief 0.5s delay. Each trial was followed by a 2-s reward-consumption period and a 1-s inter-trial interval (ITI). On the first day, a small amount of reward was also placed on the screen at the beginning of the session to motivate marmosets to touch the screen.

Once criteria were reached, marmosets proceeded to location-varying touch training: they learned to touch a black-and-white geometric ‘flower’ pattern, which appeared at middle, left or right of the screen pseudorandomly. Lastly, with the same design as the previous step, marmosets learned to refrain from touching the screen during 2-s ITIs, with each erroneous touch lengthening the ITI by 0.1 seconds.

#### Simple discrimination

This step in training was conducted for shape and colour in separate 90-trial daily sessions, and the criteria was set to 8 correct choices in 10 trials in 3 consecutive 10-trial blocks. This training served to inform marmosets that each shape or colour could be rewarding or not, depending on the session. One marmoset from each cage was randomly assigned to start with shape discrimination, while the other from the cage started with colour discrimination. For shape discrimination, in the first sessions, marmosets were presented with 2 of the 3 shapes (star, square and heart) in black at pseudorandom left vs right locations (**Figure 1**). Initially, choosing the wrong choice was counted as an error but did not terminate the trial, allowing for ‘correction’. Once the criteria were met, 1-2 additional sessions were given, in which wrong choices ended the trial with no reward. On the next set of sessions, the previously incorrect shape was now the target, and the third shape now served as the foil. At the last step of shape training, the previous foil became the target and the initial target now served as the foil. Colour discrimination followed the same process and required them to choose from 2 circular colour patches (**Figure 1A**). The colours used were red, yellow and blue. For different animals, the first target shape or colour differed, so while they could observe from their cage mate’s training, we ensured that they would never work with the same stimuli pair on the same day.

### Feature-rule switching task (FRST)

In the full version of the Feature-Rule Switching task (FRST), the colors and shapes were combined into compound stimuli for the first time (i.e. coloured shapes). The rule was to find and focus on the single target feature that was rewarded (e.g. heart or yellow) until the contingency changes (**Figure 1B, C**). On each trial (**Figure 1C**), two compound stimuli were presented on the left and right of the screen. Animals must touch the stimulus containing the target feature to obtain a reward. Incorrect choices ended the trial with no reward; and the trial was repeated until the animal made the correct choice.

A rule was considered learned once the animal reached 80% correct over the most recent 10 trials, with at least 20 trials completed under that rule. On reaching criterion, the rewarded feature switched to a new target, either within the same dimension or across dimensions (Figure 1B). The two switch types alternated within a session, so that odd-numbered switches were of one type and even-numbered switches the other; which type occurred first was counterbalanced across daily sessions. Notably, because animals were trained on Simple Discrimination equally and separately in the Colour and Shape dimensions, and were shifting targets among known features, we do not expect them to form a cognitive or attentional ‘set’ as was the case in the IEDS task. This is an important distinction between the FRST and set-shifting tasks such as the IEDS.

Once sufficient data were collected, we repeated the FRST using a new set of shape and colour features (**Figure 1D**), omitting the Simple Discrimination. That is, marmosets were presented directly with full FRST, with the new colours and shapes already combined as compound stimuli (**Figure 1D**). Data are openly accessible at: https://doi.org/10.5281/zenodo.22750042

Statistical analysis of performance data was conducted using Statistica (RRID:SCR_014213). For each animal, changes in response accuracy before and after switches (within-group effect) across conditions (between-group effect) were tested using mixed-model ANOVA (**Figure 4,5**). Additionally, the effect of accuracy of the response and that of the previous response on RT were tested using 2-way ANOVA (**Figure 6**). Post hoc Tukey’s test was conducted in case of a significant main or interaction effect.

### Reinforcement Learning Modelling

To capture trial-by-trial learning and attentional dynamics in task performance, we implemented three reinforcement learning models originally described by Talwar et al. (2024): Feature Reinforcement Learning (fRL), Combined Attention-Modulated Feature Reinforcement Learning (Ca-fRL), and Separate Attention-Modulated Feature Reinforcement Learning (Sa-fRL). The Python code (RRID:SCR_024202) was obtained with written permission from the authors from https://github.com/AnahitaTalwar/cantab-ied-models. Each model computes stimulus values based on their features, then uses a softmax function to produce the choice and updates feature and attention weights via prediction errors. We adapted the algorithm of Talwar et al. (2024) to reflect small differences in task structure without changing the underlying mathematics and applied maximum-likelihood estimation to obtain parameter estimates. Our code is openly accessible on GitHub at https://github.com/cogneurophys/Marmoset-FRST/.

#### Feature Reinforcement Learning (fRL)

The simplest model (fRL) assumes that the value of a compound stimulus, S, is the sum of its feature weights:

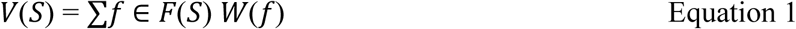

Where F(S) is the set of two features (one colour, one shape). Choices among the two on-screen stimuli were governed by a softmax rule:

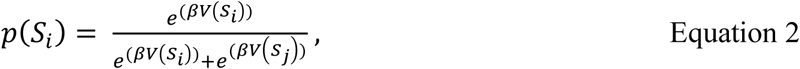

where β (choice determinism), the inverse temperature parameter controlling exploitative versus exploratory choices.

Following each trial, *t*, the weights of the chosen stimulus features are updated via

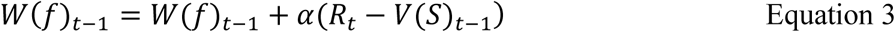

where α is the learning rate and R_t_ is the feedback or outcome of their response (correct = 1, incorrect = 0).

In this model, each individual feature (e.g., a specific colour or shape) is associated with a scalar value. After selecting a stimulus composed of two features f_1_ and f_2_, the prediction error is computed as:

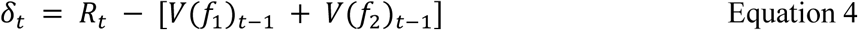

The feature values are then updated accordingly to this prediction error:

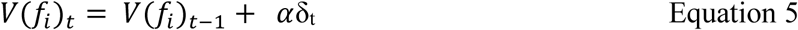

This model does not incorporate dimensional structure or attentional modulation; all features are updated equally regardless of their dimension.

#### Combined Attention-Modulated Feature Reinforcement Learning (Ca-fRL)

The Ca-fRL model extends the fRL framework by introducing attentional weighting across stimulus dimensions. In this model, stimulus value is computed as a weighted sum of its colour and shape features, with attention determining the relative contribution of each dimension. Only features belonging to the currently attended dimension are updated on a given trial, capturing selective attention to the rewarded dimension.

The attention weights for colour and shape on trial *t* is defined as:

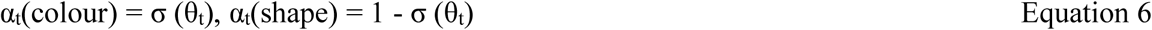

where σ denotes the sigmoid function and θ_t_ represents the attentional bias towards one dimension. The stimulus value is then given by:

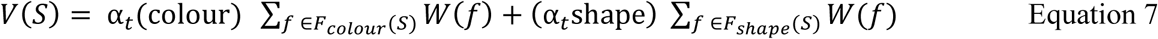

The initial attentional bias is determined by the dimension primacy parameter θ_0,_ with initial attention weights defined accordingly. On each trial, both feature values and the attentional parameter θ are updated via backpropagation of the squared-error loss using a shared learning rate α. This allows attentional weights to dynamically shift toward the dimension that best predicts reward over time.

#### Separate Attention-Modulated Feature Reinforcement Learning (Sa-fRL)

The Sa-fRL further extends the Ca-fRL framework by introducing a second learning rate, ε parameter, which is a separate learning rate that governs the updates to the dimension attentional weights, while α remains the learning rate for the feature weights. Talwar et al, (2024) describe Sa-fRL as identical to Ca-fRL except that, instead of using one α to update both feature and attention weights, dimension, θ, weights are updated with their own learning rate, ε:

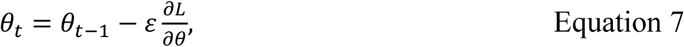

whereas feature weight still updates via

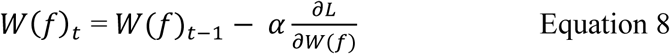

This separation allows attentional dynamics and feature value learning to operate separately, with ε indexing how rapidly attention reallocates across dimensions independently of feature learning.

#### Choice-perseveration control models

To test whether the advantage of the attention-augmented model reflects dimensional structure specifically, rather than a general tendency to repeat recent choices, we fit a perseveration control matched to Ca-fRL in parameter count. A third free parameter could improve fit simply by capturing choice stickiness — the tendency to re-select recently chosen features regardless of reward — instead of dimensional attention. A count-matched competitor lets the model comparison adjudicate between these accounts directly.

The perseveration model inherits the feature-value learning and softmax choice rule of the feature-only model without change (Eqs. X–Y); features acquire value through the same reward-driven update, and the value equations are untouched. The model adds one mechanism: a feature-level choice trace that biases the upcoming decision toward recently chosen features, independently of reward. For each feature *f* we maintain a trace κ_t(*f*), and the decision variable for option *o* becomes

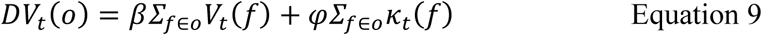

with choice probabilities given by the softmax over the two options. Here V_t(*f*) is the learned feature value, β the inverse temperature, and φ a free perseveration weight. We kept φ as a separate additive weight rather than scaling the trace by β, so that value and perseveration may operate on different scales; folding the trace under β would tie the two terms to a common gain and bias the comparison in favour of Ca-fRL. Because the trace enters the decision variable and never the value update, it captures perseveration — choice autocorrelation independent of outcome — as distinct from learning.

We considered two forms of the trace. In the primary, last-trial form, κ_t(*f*) = 1 if feature *f* belonged to the option chosen on the preceding trial and 0 otherwise; this adds a single parameter (φ) to the feature-only model, giving three free parameters {α, β, φ} — the same number as CaFRL. In a second, decaying form, the trace integrates choices over multiple trials through a delta rule,

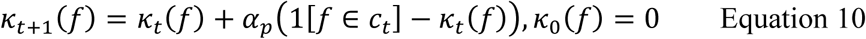

where c_t is the option chosen on trial *t* and α_p ∈ (0,1) is the trace update rate, adding a fourth parameter {α, β, φ, α_p}. The two forms are nested: fixing α_p = 1 recovers the last-trial model. We treated the three-parameter form as the primary control and the four-parameter form as a robustness check on whether perseveration extending beyond the preceding trial alters the comparison.

We fit both models to each animal’s choices by maximum likelihood, using the same optimization procedure as the other models (α and α_p constrained to (0,1), β to positive values, φ unconstrained). Models were compared by the Bayesian information criterion, BIC = −2 log *L* + *k* ln *N*, with *k* the number of free parameters and *N* the number of choice trials. Because the three-parameter perseveration model and CaFRL share the same *k*, their BIC penalties are identical, so any BIC difference between them reflects goodness of fit alone — a direct test of whether the third parameter is better spent on dimensional attention or on choice perseveration.

### Model comparison

To compare the three models while accounting for their differing numbers of free parameters, we computed the Bayesian Information Criterion (BIC) from the maximized log-likelihood for each model, animal, and stimulus set, with lower values indicating a better balance of goodness-of-fit and parsimony. As a complementary, qualitative posterior-predictive check, we also examined the correspondence between observed and model-predicted trial-by-trial error counts. Because all three models share a common feature-learning core and differ only in the presence and parameterisation of the dimensional-attention layer, BIC indexes whether dimensional attention earns its added parameters relative to feature-only learning; it does not, however, adjudicate between dimensional attention and equally flexible non-dimensional mechanisms, an important distinction we return to in the Discussion.

### Cross-species parameter comparison

To assess whether individual animals’ Ca-fRL parameters fell within the range of human values reported by Talwar et al (2024, Table 1), we compared each parameter to the human mean ± one standard deviation (learning rate 0.92 ± 0.14; choice determinism 1.34 ± 0.43; dimension primacy 1.78 ± 0.97). For learning rate, we used two one-sided tests of equivalence (TOST). Consistent with the case-study logic of primate electrophysiology, which characterises how an individual brain can operate rather than estimating a species mean, we treated each animal as a separate case and each session as a repeated measurement, running TOST within animal on the session-level estimates, separately for the original and new feature sets; a parameter was judged within the human range when both one-sided tests rejected at α = 0.05, equivalently when the 90% confidence interval of the difference lay within the bounds. Choice determinism was not equivalence-tested, because the inverse-temperature parameter is scale-dependent and not comparable across tasks. For dimensional weighting we summarised each session as commitment, |σ(θ_0_) − 0.5|, the logistic sigmoid function of the θ_0_ which has values from 0 (Shape) to 1 (Colour), adjusted by 0.5 to have a range from –0.5 to 0.5, and used as an absolute value. Thus, this commitment is the distance of the fitted weighting from equal weighting of the two dimensions. Because commitment is invariant to which dimension is favoured, it can be compared across species even though the signed primacy parameter is anchored to a different dimension in each task, and the human commitment range was obtained by passing the human dimension-primacy mean ± 1 SD through this transform. The signed per-session primacy estimates were too weakly identified for equivalence testing and are not interpreted directionally.

## ACKNOWLEDGEMENT

We are grateful to Anahita Talwar and Jonathan P. Roiser for helpful correspondence and for generously sharing anonymized subject-level parameter estimates.

## Notes

### Competing Interest Statement

The authors have declared no competing interest.

### Summary of Updates

There is no revision in the manuscript file per se. It remains identical to the previous version (v2). In v2, the bioRxiv PDF was not properly built from the uploaded PDF, so that in one section, 8 lines of text were distributed across 8 pages. This was never the case in our original PDF. I was told by bioRxiv team that this has to involve a new upload, thus I am uploading exactly the same PDF as I did for v2.

### Funding

- Canada Research Chairs: 2021-00339
- Natural Sciences and Engineering Research Council: RGPIN-2023-05079
- Canada First Research Excellence Fund: VISTA, Connected Minds

